# A GT-seq panel for walleye (*Sander vitreus*) provides a generalized workflow for efficient development and implementation of amplicon panels in non-model organisms

**DOI:** 10.1101/2020.02.13.948331

**Authors:** Matthew L. Bootsma, Kristen M. Gruenthal, Garrett J. McKinney, Levi Simmons, Loren Miller, Greg G. Sass, Wesley A. Larson

## Abstract

Targeted amplicon sequencing methods, such as genotyping-in-thousands by sequencing (GT-seq), facilitate rapid, accurate, and cost-effective analysis of hundreds of genetic loci in thousands of individuals, but studies describing detailed workflows of GTseq panel development are rare. Here, we develop a dual-purpose GT-seq panel for walleye (*Sander vitreus*) and discuss trade-offs associated with different development and genotyping approaches. Our GT-seq panel was developed using restriction site-associated DNA data from 954 individuals sampled from 23 populations in Minnesota and Wisconsin, USA. We then conducted simulations to test the utility of loci for parentage analysis and genetic stock identification and designed 600 primer pairs to maximize joint accuracy for these analyses. We conducted three rounds of primer optimization to remove loci that overamplified and our final panel consisted of 436 loci. Optimization focused on reducing variation in amplification rate among loci and minimizing the proportion of off-target sequence, both of which are important considerations for developing large GT-seq panels. We also explored different approaches for DNA extraction, multiplexed polymerase chain reaction (PCR) amplification, and cleanup steps during the GT-seq process and discovered the following: (1) inexpensive Chelex extractions performed well for genotyping, (2) the exonuclease I and shrimp alkaline phosphatase (ExoSAP) procedure included in some current protocols did not improve results substantially and was likely unnecessary, and (3) it was possible to PCR amplify panels separately and combine them prior to adapter ligation. Well-optimized GT-seq panels are valuable resources for conservation genetics and our findings should aid in their construction in myriad taxa.

## Introduction

The development of genotyping-by-sequencing (GBS) methods has allowed collection of data from thousands of markers across a genome, enabling research that was not possible using traditional genetic approaches (Davey et al., 2011; Narum et al., 2013). For example, studies using thousands of markers genotyped with restriction site-associated DNA (RAD) sequencing have shown improved sensitivity for detecting inbreeding depression (Hoffman et al., 2014), increased resolution for determining complex phylogenies (Wagner et al., 2013), and allowed researchers to observe selection on introduced alleles (Bay et al., 2019). Many genetic analyses, however, can be conducted efficiently with genotypes from tens to hundreds of single nucleotide polymorphisms (SNPs) (Anderson & Garza, 2006), making more expensive approaches such as RAD-seq unnecessary (Meek & Larson, 2019). Two such analyses that have been widely used in conservation genetics and molecular ecology for decades, are parentage analysis and genetic stock identification (GSI).

Parentage analysis involves assigning offspring to putative parents by comparing genotypes at multiple loci, while GSI infers the natal origins of individuals by leveraging baseline allele frequency estimates from populations or reporting groups. These techniques were first conducted using allozyme markers genotyped with protein electrophoresis. Although these analyses were groundbreaking, they often lacked statistical power except in cases of highly diverged stocks or simple pedigrees. The adoption of highly variable microsatellite markers in the 1990s greatly increased statistical power, allowing these two techniques to become widely adopted (Luikart & England, 1999). Despite the advances made possible by microsatellites, problems associated with homoplasy (Garza & Freimer, 1996), locus discovery (Navajas et al., 1998), and reproducibility among laboratories led researchers to explore the potential of biallelic SNPs for GSI and parentage analysis (Seeb et al., 2011).

Although SNPs are less powerful than microsatellites on a per marker basis, SNPs are more abundant in the genome, generally have low genotyping error rates, and can be genotyped using SNP panels capable of efficiently screening a large number of samples (Brumfield et al., 2003; Morin et al., 2004). Early SNP panels were constrained, however, in the availability of molecular markers suitable for genotyping and genotyping costs associated with 5’ exonuclease chemistry (Seeb et al., 2011). These constraints were significantly lessened with the proliferation of next-generation sequencing (NGS) technology. For example, methods such as RADseq facilitate quick and affordable discovery of thousands of candidate loci, which can then be selected among for specific purposes.

As SNP discovery has become less prohibitive, methods of selecting the most informative SNPs for a given study have advanced (Storer et al., 2012). Previous research has shown that information content will vary among SNPs depending on the context within which they are applied and location within the genome (i.e. coding or non-coding regions). For example, Ackerman et al. (2011) found that SNPs under diversifying selection provide increased accuracy and precision in GSI of sockeye salmon (*Oncorhynchus nerka*) from the Copper River, Alaska. In general, previous studies have shown that GSI accuracy is generally positively correlated with differentiation (e.g., *F*_ST_) and, to a lesser extent, diversity (e.g., heterozygosity) (Ackerman et al., 2011; Bradbury et al., 2011; Storer et al., 2012). Studies of SNP selection methods for parentage analysis, however, have found that high diversity is the most important attribute to consider when creating a panel (Baetscher et al., 2018). More recently, analytical techniques have shifted towards consideration of closely linked SNPs (i.e. microhaplotypes), which effectively increases the diversity at a locus and has proven useful for parentage and GSI tests (Baetscher et al., 2018; McKinney, Seeb, et al., 2017; Reid et al., 2019). While obtaining microhaplotypes using previous 5’ exonuclease methods would require independent assays for each SNP at a locus and statistical phasing, NGS technology has enabled the joint genotyping of multiple SNPs within single reads, making microhaplotype data easily obtainable through a simple modification in analytical approach.

One recently developed GBS method that improves upon previous high-throughput genotyping technologies, such as 5’ exonuclease chemistry, is Genotyping-in-Thousands by sequencing (GT-seq). This method enables genotyping hundreds of SNPs in thousands of individuals on a single NGS lane through the use of highly-multiplexed polymerase chain reaction (PCR) (Campbell et al., 2015). GT-seq does not require an allele-specific probe, can genotype multiple SNPs within an amplicon using a single primer pair, and is substantially less expensive than 5’ exonuclease chemistry, especially in the context of genotyping thousands of individuals.

Despite its benefits, GT-seq is not yet widely used outside of salmonids. Early applications to non-model organisms, however, have shown great promise for this method’s versatility, including the ability to reveal dispersal and mating patterns in a complex environment (Baetscher et al., 2019), provide insight to the ecological and evolutionary dynamics of secondary contact (Reid et al., 2019), and understand population diversity in systems that are heavily influenced by climate change (Pavinato et al., 2019). Pedigree analysis in wild populations is highly dependent upon the ability to genotype large sample sizes to increase the likelihood of detecting kin relationships, toward which GT-seq is ideally suited. Moreover, GT-seq has proven capable of generating high-quality genotypes from low-quality DNA samples (Natesh et al., 2019; Schmidt et al., 2019), making it a viable approach for monitoring endangered or elusive species.

While GT-seq panels have been developed to maximize accuracy for GSI (McKinney et al., 2019) or parentage (Baetscher et al., 2018) analyses, the potential for developing dual-purpose panels is largely unexplored. Moreover, developing GT-seq panels is a relatively involved task and, to this point, there are limited resources providing standardized workflows and guidelines for efficient panel construction (but see Campbell et al., 2015; McKinney et al., 2019). At a basic level, panel construction involves SNP discovery, SNPs selection, primer design, and panel optimization (see Baetscher et al., 2018; McKinney et al., 2019; Schmidt et al., 2019); however, within this general framework there are many decision points in panel development related to primer selection, multiplexing approaches, laboratory protocols, and analysis parameters that have yet to be addressed. We used walleye (*Sander vitreus*) from Minnesota and Wisconsin, USA, as a test case to investigate various tradeoffs associated with GT-seq panel development and optimization and leveraged our collective experience to provide guidelines for researchers developing GT-seq panels.

Walleye are an apex predator and one of the most prized sportfish throughout their native and introduced range. Recently, many walleye populations have declined across the Midwestern United States (Embke et al., 2019; Hansen et al., 2015; Rypel et al., 2018), prompting increases in stocking efforts relative to already large and long-term regional stocking programs that have existed for decades. Genetic studies have been used to guide these efforts by informing broodstock selection and general stocking practices. Genetic variation in walleye from this region was first characterized by Fields et al. (1997), who found geographic-based patterns of genetic structure, but limitations related to sample size and molecular marker choice resulted in the use of contemporary watershed boundaries as genetic management units. This research was later expanded upon by Hammen and Sloss (2019), who attempted to further define genetic structure in the Ceded Territory of Wisconsin, approximately the northern third of the state, and test whether significant genetic structure existed between distinct hydrological basins within this region. Once again, constraints associated with available molecular markers used in a system with not only low differentiation, but also extensive stocking precluded definition of fine-scale structure. This system provides an excellent model for applying genomic techniques to discriminate populations and evaluate hatchery programs using parentage analysis.

Like many intricacies of genomics research, GT-seq panel development is a process that is at once broadly generalizable to non-model organisms and highly specific to the taxa it is applied to. While the overarching steps (Fig. 1) will remain constant, there are many decision points within that will require informed thought and decision. Using walleye, a species with few well-established genomic resources, as a model, we examined the methods inherent to GT-seq panel development in a manner that identifies critical decision points in the process and illuminates the nuances associated with them. Our overarching goal was to design a dual-purpose GT-seq panel optimized for parentage analysis and GSI in walleye. The creation of this panel allowed us to address the following specific objectives: (1) investigate the tradeoffs between choosing markers for parentage analysis versus GSI, (2) explore the most efficient way to design an optimized panel, and (3) evaluate various laboratory approaches to maximizing the efficiency of GT-seq genotyping. We provide an in-depth discussion of our experiences designing the panel and outline important topics that should aid researchers in designing future GT-seq panels.

**Figure 1.**
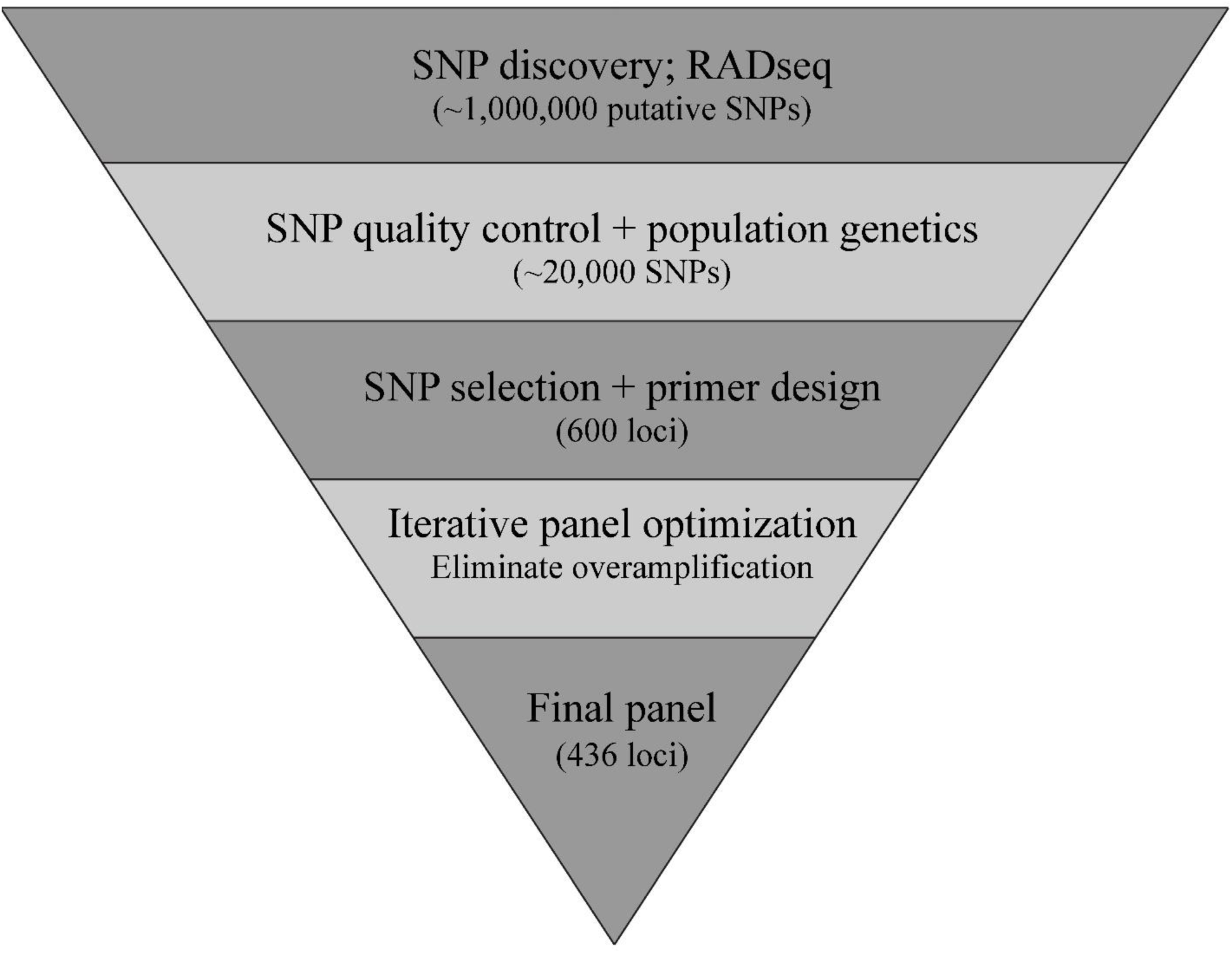
Generalized workflow describing major steps inherent to *de novo* construction of a high-density SNP panel for walleye *Sander vitreus* in Wisconsin and Minnesota, USA. Numbers of SNPs or loci present in each phase for this panel shown in parentheses.

## Materials and Methods

### Sample collection

Tissue samples were collected from adult walleye from 23 inland lakes across Wisconsin, Minnesota, and the St. Louis River (border water) (Fig. 2a, Table 1) and stored in 95% ethanol until DNA extraction. We obtained samples from as many major drainages as possible across the two states, with an emphasis on the Wisconsin and Chippewa River drainages in Wisconsin, which were difficult to differentiate using microsatellites (Hammen & Sloss, 2019); in Minnesota, sampling focused primarily on major sources of wild broodstock for stocking programs. Samples were collected by the Wisconsin and Minnesota Departments of Natural Resources using fyke nets or electrofishing. Sampling took place during the spring spawning runs of April 2015 and 2017 and fall surveys in August and September of 2015 and 2017. Stocked individuals may be tagged, or fin clipped; we inspected all sampled individuals for tags or fin clips to avoid as many individuals as possible that were of stocked origin as possible.

**Figure 2.**
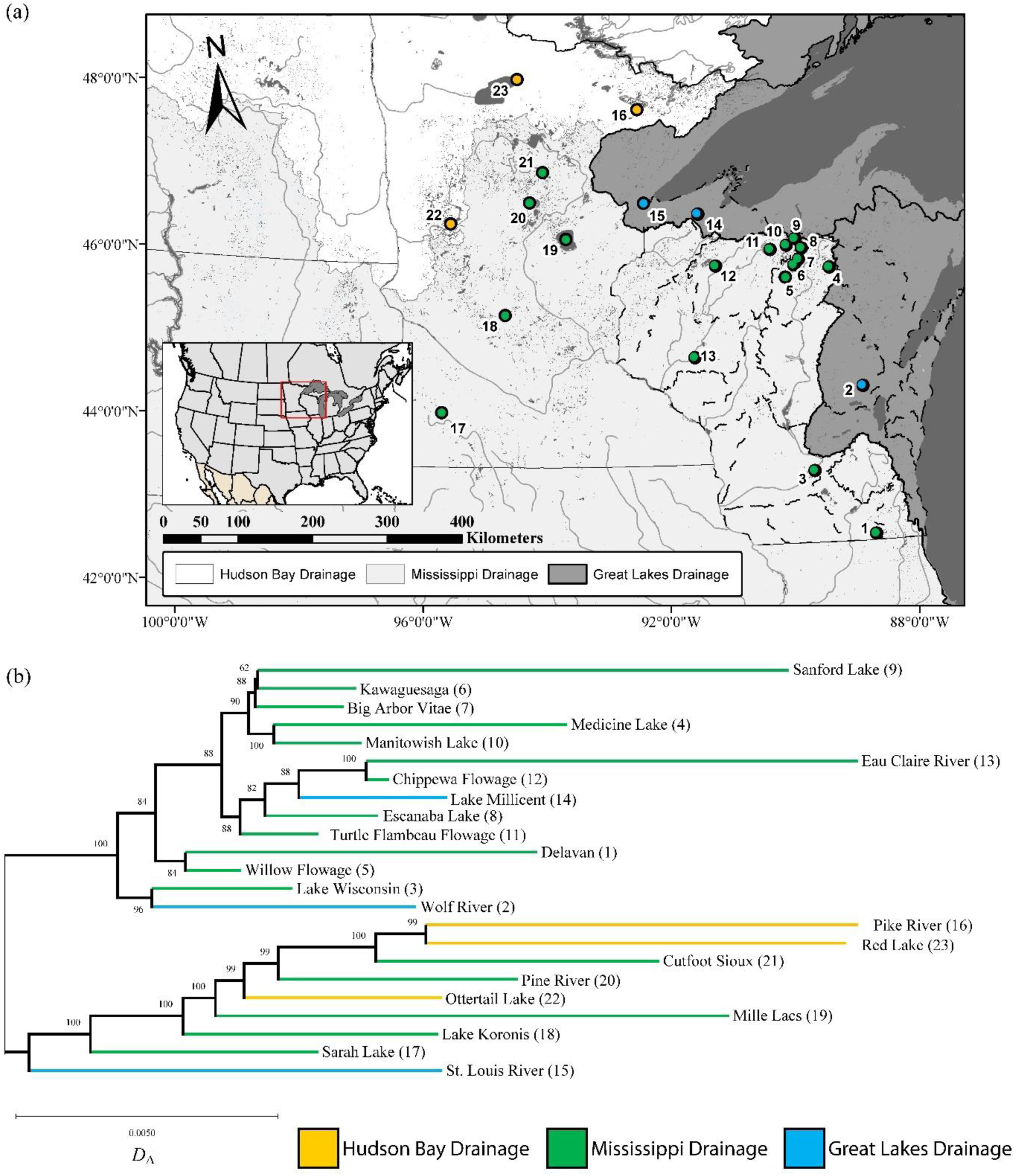
(a) Map of walleye *Sander vitreus* in Wisconsin (populations 1-14), the St. Louis River (population 15), and Minnesota (populations 16-23), USA, collection locations and (b) dendrogram of sampled populations with bootstrap support (n = 1000) estimates above nodes. Branch lengths correspond to genetic distances estimated using Nei’s *D*_A_. Figures color coded according to major drainage of origin (Hudson Bay: yellow, Mississippi: green, Great Lakes: blue) and numbered with respect to order in Table 1.

**Table 1.**
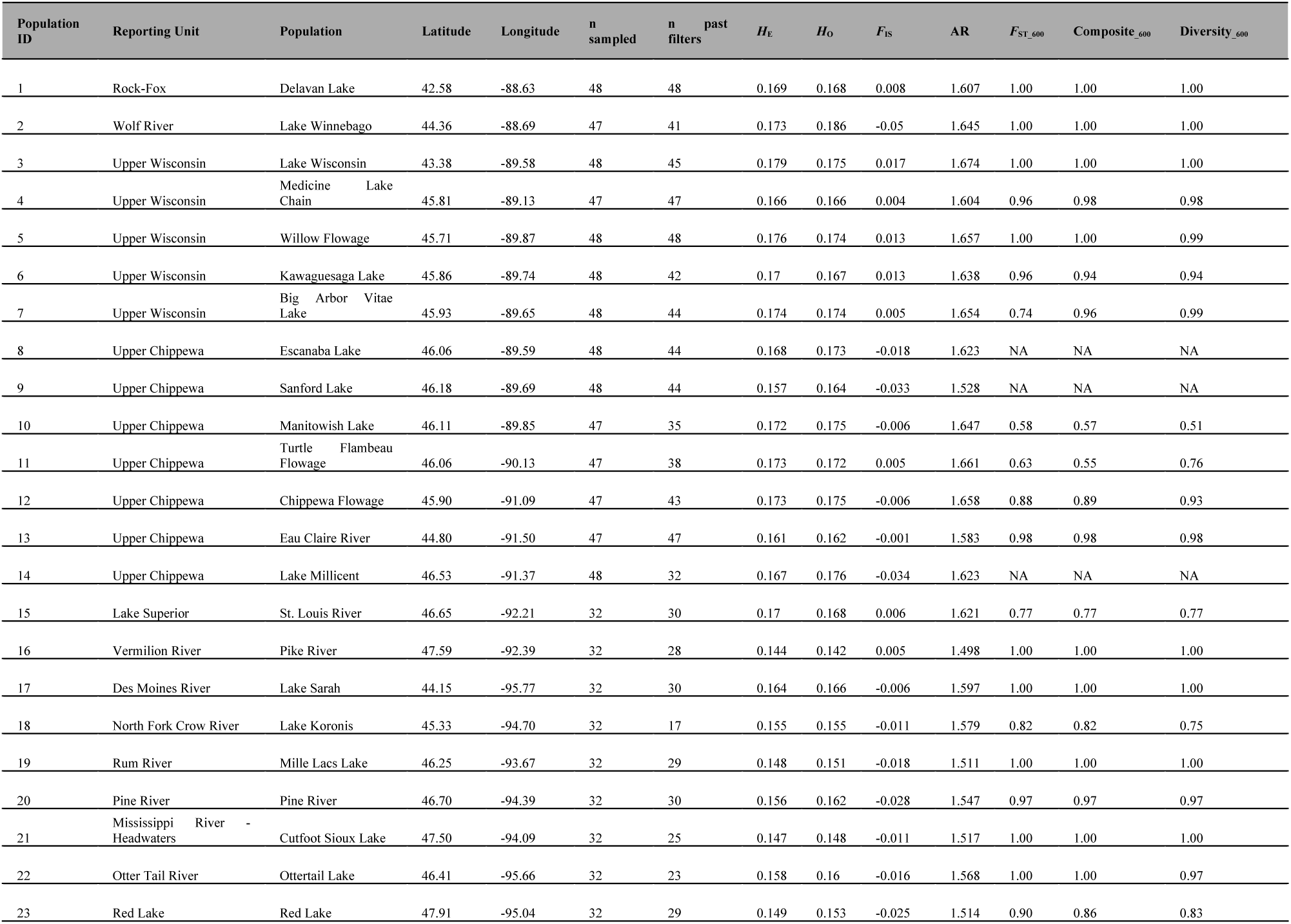
Information on walleye *Sander vitreus* collections from 23 sites in Wisconsin and Minnesota. Reporting units are aggregations of genetically similar populations grouped for GSI analysis, n past filters is the number of individuals missing genotypes at < 20% of SNPs and retained after quality filtering. Diversity statistics calculated using 20,579 SNPs. The F_ST_600_, Composite__600_, and Diversity__600_ columns are the percent correct assignment to reporting group for each population with 100% simulations conducted using the corresponding panel.

### Preparation of RAD sequencing libraries

Genomic DNA was extracted in a 96-well format with Qiagen DNeasy Blood and Tissue Kits. Extracted DNA was quantified using a Quant-iT PicoGreen dsDNA Assay Kit (Invitrogen, Waltham, MA) and normalized to 20ng/μl. DNA was then prepared for RADseq library preparation following the BestRAD protocol (Ali et al., 2016). Briefly, DNA was digested in a 2 μl reaction with the restriction enzyme *SbfI*, and biotinylated barcode adaptors were ligated to the 5’ cut ends. DNA shearing was conducted using a 12.5 μl fragmentase reaction. Library preparation was conducted using an NEBNext Ultra DNA Library Prep Kit for Illumina (NEB, Ipswich, MA), with a 12-cycle PCR enrichment. RAD library quality was inspected on a 2% agarose gel before undergoing a final AMPure XP (Beckman Coulter, Indianapolis, IN) purification and quantification on a Qubit 2.0 Fluorometer (ThermoFisher Scientific, Waltham, MA). Libraries were sequenced using paired-end (PE) 150 technology on a HiSeq 4000 (Illumina, San Diego, CA) at the Michigan State University Genomics Core Facility or Novogene Corporation, Inc. (Davis, CA). Sequencing was conducted to achieve a target of over one million retained reads per individual.

### Analysis of RAD data to discover SNPs

Loci were identified and genotyped in STACKS v.2.2 (Rochette et al., 2019) without using gapped alignments. Raw reads were demultiplexed and barcodes were trimmed in *process_radtags* (parameter flags: -e *SbfI*, -c, -q, -filter_illumina, -r, --bestrad). RAD-tags were assembled into putative RAD loci with *ustacks* using the bounded model (bound_high = 0.05, --disable-gapped) and allowing for a maximum of three nucleotide mismatches (−M = 3) and four stacks per locus (−max_locus_stacks = 4), as well as a minimum depth of three (−m = 3). The calling of haplotypes from secondary reads was disabled (−H). A catalog of consensus loci was assembled in *cstacks* using the two individuals with the highest number of retained reads from each population, allowing a maximum of three mismatches between sample loci (n = 3, --disable-gapped). After matching all samples against the catalog in *sstacks* (--disable-gapped), data were oriented by locus with *tsv2bam*, and individual genotypes were called in *gstacks*, with paired-end reads incorporated. Genotypes were exported in variant call format (vcf) using *populations*, with loose filtering parameters (SNPs present at > 5% of individuals, minimum minor allele frequency of > 0.005).

Comprehensive filtering of individuals and genotypes was conducted in vcftools v0.1.15 (Danecek et al., 2011) by: 1) removing individuals missing > 20% of SNP calls, 2) removing SNPs that were missing in > 20% of individuals, and 3) removing SNPs that were not in the first 140 base pairs of the RAD-tag, effectively reducing the dataset to include SNPs detectable using single-read (SR) 150 sequencing to simplify downstream amplicon design; to control for genotyping error, SNPs with a minor allele count ≤ 3 were also removed. Putative duplicated loci were identified in HDplot (McKinney, Waples, et al., 2017) (H > 0.5, −7 < D < 7) and removed with vcftools. Retained individuals and SNPs were used to form whitelists for input into *populations* that output a filtered vcf of multi-SNP haplotypes, which was then filtered to remove loci with more than 10 alleles and used in simulations for locus selection. We also estimated single-SNP *F*_IS_ across all populations using diveRsity v1.9.90 (Keenan et al., 2013) and excluded any SNPs with *F*_IS_ values > 0.2 or < −0.2 from locus selection. Additionally, loci with a SNP in the first 10 base pairs of the RAD-tag were excluded to allow room for forward primer design.

### Analysis of population structure, locus selection, and panel assessment

To understand population structure in our system and ensure that selected loci could facilitate accurate parentage assignment and GSI, we evaluated patterns of genetic divergence using pairwise *F*_ST_ (Table S1) estimated in Arlequin v3.5.2 (Excoffier & Lischer, 2010) and constructed a dendrogram (Fig. 2b) using Nei’s distance in poppr v2.8.2 (Kamvar, Tabim, & Grünwald, 2014). These analyses facilitated identification of population pairs that would be challenging to discriminate and supported historical data suggesting several populations were founded from hatchery sources located outside of their drainage basin (Escanaba Lake, Sanford Lake, and Lake Millicent in Wisconsin); these populations were removed from simulations of panel accuracy to ensure that selected loci would best represent the natural genetic patterns of the region.

After initial population genetic analyses, loci were selected for primer development by constructing several test panels from the RAD data and simulating assignment accuracy for parentage and GSI. Previous research suggested that choosing loci with greater genetic differentiation (e.g., *F*_ST_) should maximize accuracy for GSI (Ackerman et al., 2011; Storer et al., 2012), while choosing loci with higher diversity (e.g., heterozygosity and number of alleles) maximizes accuracy for parentage (Baetscher et al., 2018). We therefore constructed the test panels using single-SNP *F*_ST_ estimated in diveRsity v1.9.90 (Keenan et al., 2013) as well as expected heterozygosity at a multi-SNP haplotype (*H*_E_mhap_) and the number of alleles at a locus estimated in adegenet v2.1.1 (Jombart & Ahmed, 2011). All simulations were conducted with genotypes coded as multi-SNP haplotypes.

GSI accuracy for each panel was assessed via 100% simulations implemented in rubias (Moran & Anderson, 2018) using the *assess_reference_loo* function (mixsize = 200, reps = 1000). Populations were aggregated into reporting units based on hydrological basins (Table 1). Collections within a simulation were drawn from a Dirichlet distribution with all parameters equal to 10 (i.e., each simulation’s prior contained approximately equal proportions of each population for the given reporting unit). Individuals were assigned to reporting groups if they had a cumulative probability of > 70%. Unfortunately, limited sample sizes in some reporting units prevented creation of separate training and holdout datasets as suggested by Anderson (2010), thus assignment accuracies presented here may be upwardly biased and would need to be reassessed more thoroughly for populations involved in an applied study.

Parentage simulations were run in CKMRsim (Anderson, https://zenodo.org/record/820162), which employs a variant of the importance-sampling algorithm of Anderson and Garza (2006) that allows for more accurate estimates of very small false-positive rate (FPR: per-pair rate of truly unrelated individuals being inferred as related) relative to those obtained using standard Monte Carlo methods (Baetscher et al., 2018). Parentage analyses were conducted following the methods of Baetscher et al. (2018), whereby log-likelihood ratios between a tested relationship and the hypothesis of no relationship are computed from the calculated probabilities of genotype pairs for related individuals simulated from allele frequency estimates. Distributions of simulated log-likelihood ratios are then used to compute FPRs. Using this approach, we estimated FPRs for parent-offspring (PO), full-sibling (FS), and half-sibling (HS) relationships at false-negative rates (FNR: per-pair rate of truly related individuals being inferred as unrelated) ranging from 0.01 to 0.1.

Panels of 600 unique loci were iteratively selected, choosing loci based first on rank *F*_ST_ then rank *H*_E_mhap_, and their utility was tested by conducting GSI tests and parentage simulations. We ultimately defined three panels of 600 loci that best described the tradeoffs between markers selected based on *F*_ST_ and heterozygosity. Loci in these panels were chosen by selecting 1) the top 600 loci based on *F*_ST_, 2) the top 300 loci based on *F*_ST_ and 300 based on *H*_E_mhap_, and 3) the top 600 loci based on *H*_E_mhap_. These panels are hereafter referred to as *F*_ST_600_, Composite__600_, and Diversity__600_, respectively. Through further testing, we determined that a variation of the Composite__600_ panel, with 250 loci based on *H*_E_mhap_ and 350 loci based on *F*_ST_, delivered optimal performance for GSI and parentage analyses and proceeded to design primers for the selected loci.

### Primer Design

To design PCR primers for the selected loci, their consensus sequences were subset from the STACKS catalog into a FASTA file for import into Geneious Prime® 2019.1.1 (https://www.geneious.com). The vcf file produced in the vcftools step containing all SNPs and alleles within a consensus sequence was included to ensure primers were properly designed (i.e., should a SNP fall within a primer binding region, a degenerate nucleotide could be inserted or the primer re-designed). Primer pairs were iteratively designed, with optimal target parameters defined as a primer length of 20 bp, product size of 140 bp to facilitate genotyping with SR chemistry, T_m_ of 60° C, GC content of 50%, and no more than four of the same base repeated consecutively (i.e., poly-X repeats). Primers identified as matching one or more off-target sites, which could lead to amplification of multiple products, were redesigned. Given that not all 600 candidate loci initially identified were suitable candidates for primer development, we continued to iteratively select loci and design associated primers until we reached our target of 600 loci. Unfortunately, the loci selected for primer design were based on data containing a subset of individuals with discordant encoded and true identities as a result of transposition of barcodes during demultiplexing. Despite these discrepancies, the effect was likely minor as only 8% of individuals were incorrectly assigned to reporting units prior to simulation. Simulation results shown here were conducted using corrected data.

### GT-seq optimization

GT-seq was conducted following the methods of Campbell et al. (2015), with modification to the multiplex thermal cycling conditions (95 °C hold for 15 min; five cycles of 95 °C for 30 s, 5% ramp to 57 °C for 2 min, 72 °C 30 s; and 10 cycles of 95 °C for 30 s, 65 °C for 30 s, and 72 °C 30 s) and post-normalization dual-sided SPRI size-selection and purification (0.6X plus 0.4X) to further restrict the product size range (e.g., primarily toward removal of primer inter-hybridization). Final library quality control consisted of confirmation of amplification and barcoding by SYBR Green-based RT-qPCR (Stratagene Mx3005P QPCR System, Agilent, Santa Clara, CA), visualization on a 2% agarose E-Gel (Invitrogen, Carlsbad, CA), and quantification using picogreen. Libraries were then sequenced at the University of Wisconsin-Madison Biotechnology Center (UWBC) DNA Sequencing Facility on a MiSeq (Illumina) using 2 × 150 bp flowcells.

Demultiplexed amplicon sequencing data were processed using *GTscore v1.3* (McKinney et al., 2019). *GTscore* generates *in-silico* primer-probe sequences from a catalog of loci generated in STACKS, that are then matched to amplicon sequences and call genotypes for individual SNPs as well as multi-SNP haplotypes. *GTscore* also enables separation of on-target sequence reads (i.e., reads containing both an *in-silico* primer and associated probe) from reads produced as a result of primer cross-hybridization. Primer-probe file development was accomplished with *sumstatsIUBconvert.pl* by obtaining the IUB code information for each SNP from the sumstats.tsv file produced in the STACKS pipeline, converting catalog sequences produced in the STACKS pipeline to FASTA sequences using *catalog2fasta.pl*, and merging IUB code information with the catalog.fasta using *fasta2IUB.pl*. This primer-probe file was then input for *AmpliconReadCounter.pl*, along with an individual’s fastq file, to produce read count summaries of primers and probes.

Overall, we conducted three rounds of panel optimization to identify and remove loci that had disproportionately high amplification rates (i.e., “overamplifiers”) and ensure that our panel was capable of delivering a high proportion of on-target reads for each locus as well as homogeneous amplification rates among loci. The first round of optimization used DNA from a single walleye from Sanford Lake, WI, while the second and third rounds were conducted on subsets of 24 individuals from each of four populations (96 individuals total) originally included in the RADseq study: Delavan Lake, Medicine Lake, and the Wolf River in Wisconsin and the Pine River in Minnesota. Upon completing the final optimization, the characteristics of retained loci were compared to those of loci culled from the panel. This was done by performing a Welch’s two sample t-test (α = 0.05) between the GC:AC ratio of primers that were retained and those culled and between the GC:AC ratio of DNA templates retained and culled, based on the first 140 bp of the template as this was the region in which SNPs were targeted.

GT-seq libraries from each round were collectively analyzed for PCR accuracy and uniformity. Accuracy was measured by calculating the proportion of reads containing *in-silico* primer sequences (total reads) relative to those that also contained *in-silico* probes. Uniformity of amplification among loci was determined by calculating the proportion of total reads that were allocated to the top 10% of loci, based on locus read counts (prop_reads_T10); if amplification was perfectly uniform across loci, we would expect prop_reads_T10 to account for exactly 10% of total reads. Given that amplification rates vary substantially within a panel, we compared among locus performance by plotting the relative log_10_ abundance of total and on-target reads at each locus in descending order, which facilitated visual identification of overamplifiers. As among-locus amplification rates evened out after the first optimization, the on-target proportion of reads at each locus became a factor in retaining or excluding loci during the second optimization.

### Testing methodological modifications and performance analysis

During panel optimization, we compared the quality of GT-seq libraries prepared from DNA extracted with Qiagen DNeasy and a more cost-effective chelating resin-based procedure. Performance of libraries was compared using Bonferroni corrected (α = 0.016) Tukey’s HSD for the number of on-target reads and the proportion of total reads that were on-target, after determining whether significant differences existed among libraries via a one-way ANOVA (α = 0.05). DNA was extracted from the 96 test individuals twice, first using Qiagen DNeasy and again with a 10% Chelex 100 (200-400 mesh; Bio-Rad, Hercules, CA) solution containing 1% each of Nonidet P-40 and Tween 20 (Millipore Sigma, St. Louis, MO).
Additionally, we aimed to further reduce the cost per sample by evaluating the need for certain library preparation steps. Specifically, we compared results with and without the exonuclease I and shrimp alkaline phosphatase (ExoSAP) procedure included in Campbell et al. (2015) to remove PCR inhibitors and free nucleotides. GT-seq was therefore conducted on all individuals in triplicate: 1) Qiagen with ExoSAP, 2) Chelex with ExoSAP, and 3) Chelex without ExoSAP, and all tests were sequenced on the same MiSeq lane. Finally, we tested whether the number of loci that could be genotyped simultaneously could be increased by conducting multiple PCRs. We accomplished this by dividing our optimized primer panel into two non-overlapping primer pools before multiplex PCR amplification. We then merged PCR products from the separate pools prior to the barcoding PCR. The sequencing performance of this joint panel was then compared to the single multiplex containing the full panel using a Welch’s two sample t-test (α = 0.05).

We examined genotype concordance between RADseq and GT-seq across GT-seq read depths using the fully optimized panel in the third round. Genotypes were called using *PolyGen* (McKinney et al., 2018), an extension of the *GTscore* pipeline that uses the same maximum-likelihood algorithm as STACKS v1 for diploid, bi-allelic loci. Because low read depths can lead to high estimates of genotyping error, thereby increasing rates of allelic dropout (Catchen et al., 2013), genotypes were only compared if they had greater than 60× coverage in RADseq. We then modeled the relationship between GT-seq read depth and genotype concordance using only read depths with more than 30 genotypes to ensure that estimates of genotype concordance at a given depth had adequate sample sizes.

As a final proof of concept, we tested the optimized panel on a sample of 570 walleye obtained from Escanaba Lake, WI, using the methods described above to estimate the variance in read depth among loci within a pool. We retained only loci present in more than 70% of individuals and individuals genotyped at more than 70% of loci.

## Results

### Analysis of ascertainment dataset

A total of 954 individuals from 23 populations were RAD sequenced, with an average of 42 individuals per population (Table 1). Sequencing yielded 1,313,358 retained reads on average per individual (range = 8,941 - 8,176,163). Initial sequence data were used to identify 682,223 putative SNPs. After passing sequence data through quality filters, 839 individuals and 20,597 SNPs were retained (Table S2).

Population estimates of *H*_O_ (0.144 - 0.179), allelic richness (1.498 - 1.674), and *F*_IS_ (−0.050 - 0.017) were relatively similar across locations (Table 1). Populations from Minnesota had slightly lower diversity, which may be due to ascertainment bias as 14 of the 23 populations were from Wisconsin. The highest genetic differentiation was observed between populations from Minnesota and Wisconsin, with further structuring by drainage basin within each state (Fig. 2b, Table S1). Structuring was higher in Minnesota, with most populations showing a relatively high degree of isolation (average *F*_ST_ = 0.07, Table 2). Structure in Wisconsin was shallower (average *F*_ST_ = 0.03, Table 2) and only loosely correlated with drainage basins. From these results, we constructed 13 reporting groups to facilitate GSI to identifiable genetic units (Table 1). All the reporting groups from Minnesota contained single populations, whereas in Wisconsin, while the Rock-Fox and Wolf River groups contained single populations, the Wisconsin and Chippewa River groups each contained five populations. Some single populations in the Wisconsin and Chippewa Rivers were distinctly identifiable (e.g., Eau Claire River, Medicine Lake), but we grouped these populations within their drainage basin of origin as the panel will likely be used this way for management purposes.

**Table 2.** Summary of pairwise *F*_ST_ comparisons between walleye *Sander vitreus* populations grouped by state of origin. Abbreviations are Wisconsin (WI) and Minnesota (MN).

|  | WI-WI | MN-MN | WI-MN |
| --- | --- | --- | --- |
| Max | 0.106 | 0.142 | 0.142 |
| Mean | 0.032 | 0.068 | 0.072 |
| Min | 0.001 | 0.019 | 0.026 |

### Locus selection and panel assessment

GSI accuracy was similar among the three panels, with < 1% difference in average accuracy between the panel with loci chosen based solely on differentiation (*F*_ST_600_) and the panel based solely on diversity (Diversity__600_) (Fig. 3, Table 3). Average assignment accuracy was > 90% for nine of the 13 reporting units in all panels (Fig. 3a). The remaining four reporting units had average assignment accuracies ranging from 78% to 86%. Three of these units (upper Chippewa River, WI; St. Louis River, MN/WI; and Red Lake, MN) are known to have admixed stocking histories, while the fourth, North Fork Crow River, MN, included Lake Koronis, which had the fewest individuals retained after filtering (n = 15). Misassigned individuals from the St. Louis River, MN, and Red Lake, MN groups primarily assigned to the Pike River, MN, an unsurprising result given that fish from the Pike River contributed to the recovery of the collapsed walleye fishery in Red Lake (Logsdon et al., 2016) and fish in the St. Louis River watershed. Misassignments from the Upper Chippewa basin primarily assigned to the Upper Wisconsin basin due to the lower differentiation described previously.

**Figure 3.**
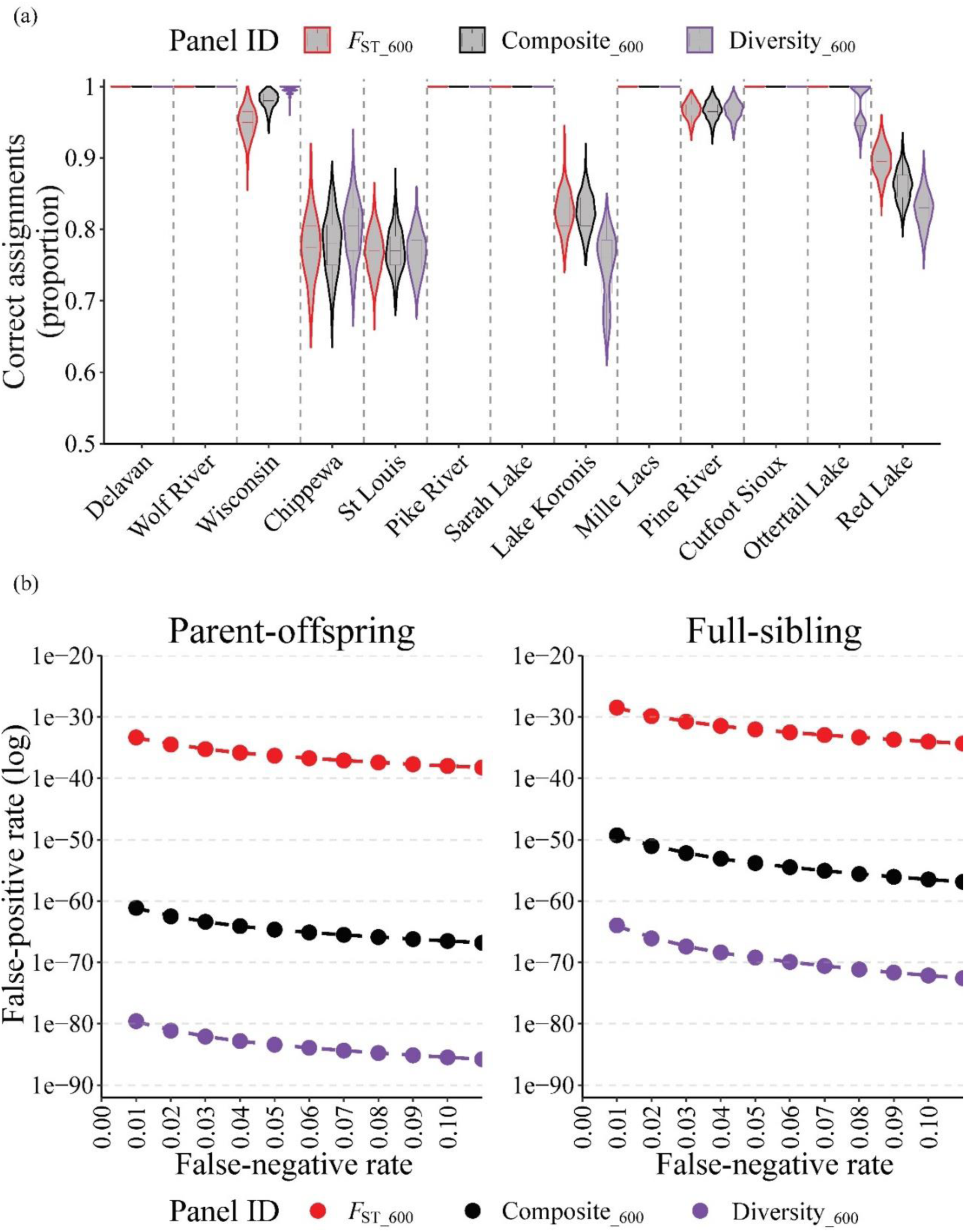
(a) Violin plots showing densitity distributions of accuracy estimates from 100% simulations of 23 populations of walleye *Sander vitreus* in Wisconsin and Minnesota, USA, performed using 1,000 iterations for each test panel by reporting unit and (b) simulated false-positive rate (FPR) estimates across a range of false-negative rates (FNR). Figures color coded according to SNP panel tested: *F*_ST_600_ (red, 600 rank *F*_ST_ loci), Composite__600_ (black, 300 rank *F*_ST_ and 300 rank *H*_E_mhap_ loci), and Diversity__600_ (purple, 600 rank *H*_E_mhap_ loci).

**Table 3.** Summary statistics by SNP panel tested for walleye *Sander vitreus* in Wisconsin and Minnesota, USA, including: average *F*_ST_, heterozygosity (*H*_E_mhap_), assignment accuracy to population and reporting unit of origin in 100% simulations, and estimated false-positive rates (FPR) for a given kin relationship at a false-negative rate (FNR) of 0.01.

| | $F_{ST\_600}$ | Composite<sub>600</sub> | Diversity<sub>600</sub> |
| --- | --- | --- | --- |
| Average $F_{ST}$ | 0.117 | 0.076 | 0.047 |
| Average $H_{E\_mhap}$ | 0.389 | 0.569 | 0.633 |
| Average accuracy by reporting unit | 0.937 | 0.937 | 0.929 |
| Average accuracy by population | 0.864 | 0.861 | 0.862 |
| Parent-offspring FPR (FNR = 0.01) | $4.68 \times 10^{-34}$ | $7.92 \times 10^{-62}$ | $2.74 \times 10^{-80}$ |
| Full-sibling FPR (FNR = 0.01) | $3.42 \times 10^{-29}$ | $5.34 \times 10^{-50}$ | $1.16 \times 10^{-64}$ |
| Half-sibling FPR (FNR = 0.01) | $6.44 \times 10^{-6}$ | $2.56 \times 10^{-10}$ | $2.06 \times 10^{-13}$ |

The populations with the lowest assignment accuracies were found in the Chippewa River and Wisconsin River reporting groups (Table S3, S4, S5), particularly in northern Wisconsin near the headwaters of the Chippewa and Wisconsin River drainages, and included Big Arbor Vitae Lake (*F*_ST_600_ accuracy = 74%), Manitowish Lake (*F*_ST_600_ accuracy = 58%), and Turtle Flambeau Flowage (*F*_ST_600_ accuracy = 63%). A large portion (> 10%) of the simulated individuals from these populations could not be assigned to any population, providing further support for the genetic similarity of these two reporting groups. A high proportion of individuals from Big Arbor Vitae Lake were assigned to Manitowish Lake (12%) and vice versa, from Manitowish Lake to Big Arbor Vitae Lake (20%). Most misassignments in the Turtle Flambeau Flowage were to Kawaguesaga Lake (16%). Populations with high misassignment rates also tended to have short branch lengths in the dendrogram and were often located near the root of a clade (Fig. 2b). Furthermore, the two populations from the upper Chippewa basin (Manitowish Lake and Turtle Flambeau Flowage) had lower pairwise *F*_ST_ values, on average, relative to populations from the upper Wisconsin basin than they did with other populations from the upper Chippewa basin.

The Diversity__600_ panel had the highest accuracy for assigning kin relationships, the Composite__600_ panel showed intermediate performance and the *F*_ST_600_ panel had the lowest accuracy rate (Fig. 3b, Table 3). For all panels, FPRs were < 10-20 for PO and FS relationships, indicating all panels would perform adequately for reconstructing most relationships in most study systems. Inter-panel performance did, however, range widely, from an FPR of 4.68 × 10-34 for *F*_ST_600_ to 2.74 × 10^−80^ for Diversity__600_ panel at an FNR of 0.01. Within panels, FPR was inversely related to FNR.

Primers were designed using a modified Composite__600_ panel, with 250 loci chosen based on *H*_E_mhap_ and 350 chosen based on *F*_ST_, as this panel delivered the best joint accuracy for GSI and kinship analyses (Fig. 3, Table 3). Of the initial 600 loci initially selected for primer design, 100 were not suitable for primer design, and thus, iterative selection of loci meeting primer design requirements was continued until the targeted number of *F*_ST_ and diversity markers was met.

### GT-seq optimization

Initial amplification and MiSeq sequencing of all 600 loci yielded 4,655,071 reads containing intact i7 barcode sequences, with 4,150,910 reads (89%) matching *in-silico* primer sequences. Locus specificity was considered via the proportion of total reads that were on-target, which was 1,031,707 (24.9%) (Table 4). In terms of amplification uniformity among loci, prop_reads_T10 accounted for 3,526,201 (85.0%) of the 4,150,910 total reads. A cutoff of 3,000 reads per locus was then visually identified (Fig. 4a); loci producing more than 3,000 reads (n = 123) were deemed overamplifiers and discarded prior to further optimization.

**Table 4.** Summary of GT-seq optimization runs for walleye *Sander vitreus* in Wisconsin and Minnesota, USA. Rows report number of primer pairs targeted, number of reads with intact i-7 barcodes (retained reads), number of retained reads with *in-silico* primer sequences (total reads), number of total reads with *in-silico* probe sequences (on-target reads), percent of total reads on-target, percent of total reads allocated to the 10% of loci tested with highest rank total read counts, average number of SNPs per locus, and average GC content in the forward and reverse primers.

|  | Test 1 | Test 2 | Test 3 |
| --- | --- | --- | --- |
| Total primer pairs | 600 | 477 | 436 |
| i7 reads | 4,655,071 | 12,653,262 | 7,282,101 |
| i7 reads w/ primers (total reads) | 4,150,910 | 9,347,591 | 6,827,424 |
| i7 reads w/ primers & probes (on- target) | 1,031,707 | 3,268,293 | 6,262,523 |
| On-target percent of total reads | 24.9% | 35.0% | 91.7% |
| Percent reads in top 10% of loci | 85.0% | 72.5% | 36.6% |
| mean SNPs per locus | 2.06 | 2.00 | 1.97 |
| mean GC percent forward primer | 51.0% | 50.4% | 50.3% |
| mean GC percent reverse primer | 49.0% | 48.3% | 48.2% |
| mean $F_{ST}$ | 0.133 | 0.133 | 0.133 |
| mean $H_{E\_mhap}$ | 0.425 | 0.415 | 0.416 |

**Figure 4.**
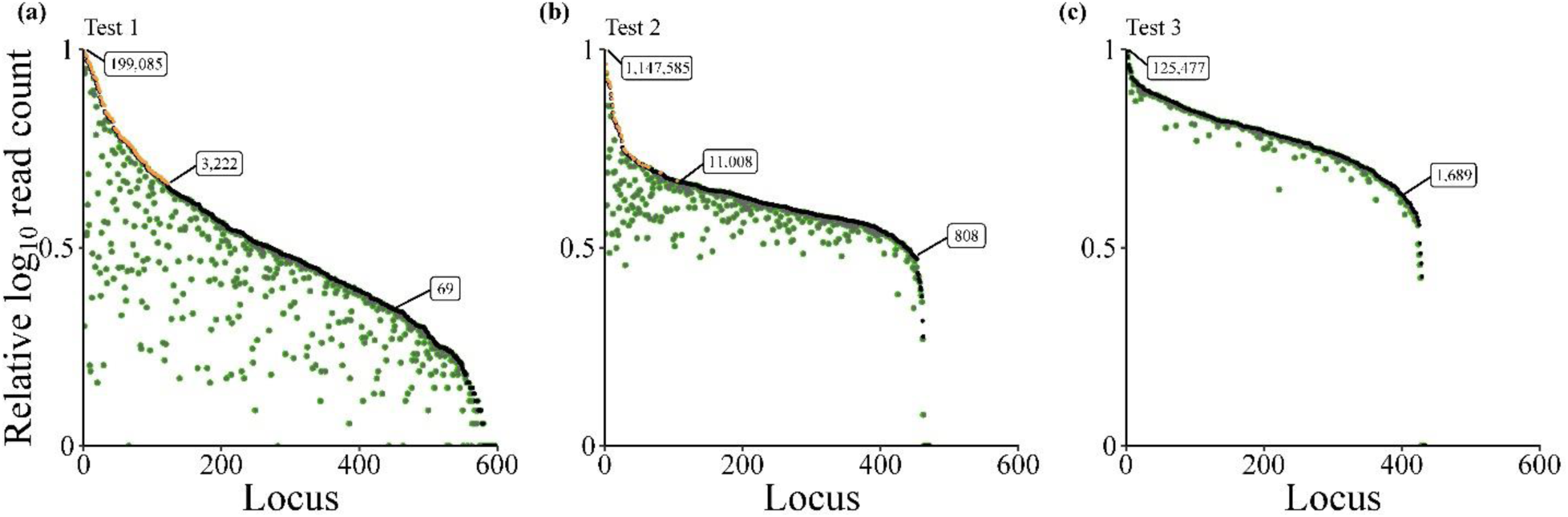
Relative log_10_ total read counts per locus (black) and relative log_10_ on-target read counts per locus (green) of the GT-seq panel for walleye *Sander vitreus* in Wisconsin and Minnesota, USA, prior to optimization (a, 600 loci), after first optimization (b, 477 loci), and after second optimization (c, 436 loci). Loci identified for culling during optimization steps shown in orange. Raw read counts annotated in boxes.

For the second round of optimization, the remaining 477 primers pairs produced 12,653,262 reads containing intact i7 barcode sequences, and 9,347,591 (74%) matched *in-silico* primer sequences. Locus specificity improved, with 3,268,293 (35.0%) of the total reads successfully aligning to *in-silico* probe sequences (Table 4). Improvement was also observed in the uniformity of amplification across loci, with prop_reads_T10 equating to 72.5% (6,776,302) of total reads. Because locus performance was less variable in this round of testing, the individual on-target proportion of reads at a locus was also considered while culling undesirable loci. As such, loci visually identified as overamplifiers were again discarded if they did not display high on-target read proportions (n = 41, Fig. 4b).

The third GT-seq test was used to determine the functional performance of the panel and aimed to target 858 SNPs across 436 loci (Fig. 4c). This test produced 7,282,101 reads with intact i7 barcodes, and 6,827,424 (94%) matched to *in-silico* primers. Locus specificity of primer pairs improved greatly in this test, as 6,262,523 (91.7%) of the total reads were also on-target (Table 4). Likewise, the variation in amplification rates across loci decreased as evidenced by prop_reads_T10 decreasing to 36.6% (2,148,932) of the total reads.

Upon completion of panel optimization, a small but significant difference was observed between the GC content of primers that were retained (mean = 49.2%) and primers that were removed (mean = 51.4%, df = 602, t = 5.4, p < 0.001). Similar differences were found when comparing the GC content of the DNA template; significantly higher GC proportions were present in templates that were culled from the panel (mean = 47.8%) than templates that were retained (mean = 45.5%, df = 359, t = 3.8, p < 0.001). Additionally, a total of 88 primer pairs in the original panel contained at least one degenerate nucleotide, 72 (81%) of which were in the forward primer. After optimization, 56 of the initial 88 (64%) were retained. In comparison, of the 512 initial primer pairs that did not have degenerate primers, 380 (74%) were retained. The average *F*_ST_ for the most informative SNP at a locus and the average *H*_E_mhap_ did not change appreciably between the initial and fully optimized panels (Table 4).

### Methodological modifications and performance analysis

Significant differences for on-target read counts and the proportion of total reads that were on-target were detected among genomic DNA extraction and purification method combinations. Subsequent analysis using Tukey’s HSD revealed that Chelex-extracted DNAs produced the highest on-target read count, and Qiagen-extracted DNAs with ExoSAP-purification produced the lowest (Fig. 5, p < 0.001). While the proportion of on-target reads did not differ between Chelex with ExoSAP and Qiagen with ExoSAP, both methods produced a significantly lower proportion of on-target reads than the Chelex-only library (Fig. 5, p < 0.001). Additionally, when comparing results from the full panel of 436 primer pairs to those obtained using the same panel divided into two unique multiplexes of 209 and 227 primer pairs (n = 436) and repooled prior to barcoding, no significant differences were found in total primer reads (df = 860, t = 0.10, p = 0.92), on-target reads (df = 858, t = 0.16, p = 0.87), or the proportion of total reads that were on target (df = 806, t = 0.66, p = 0.51).

**Figure 5.**
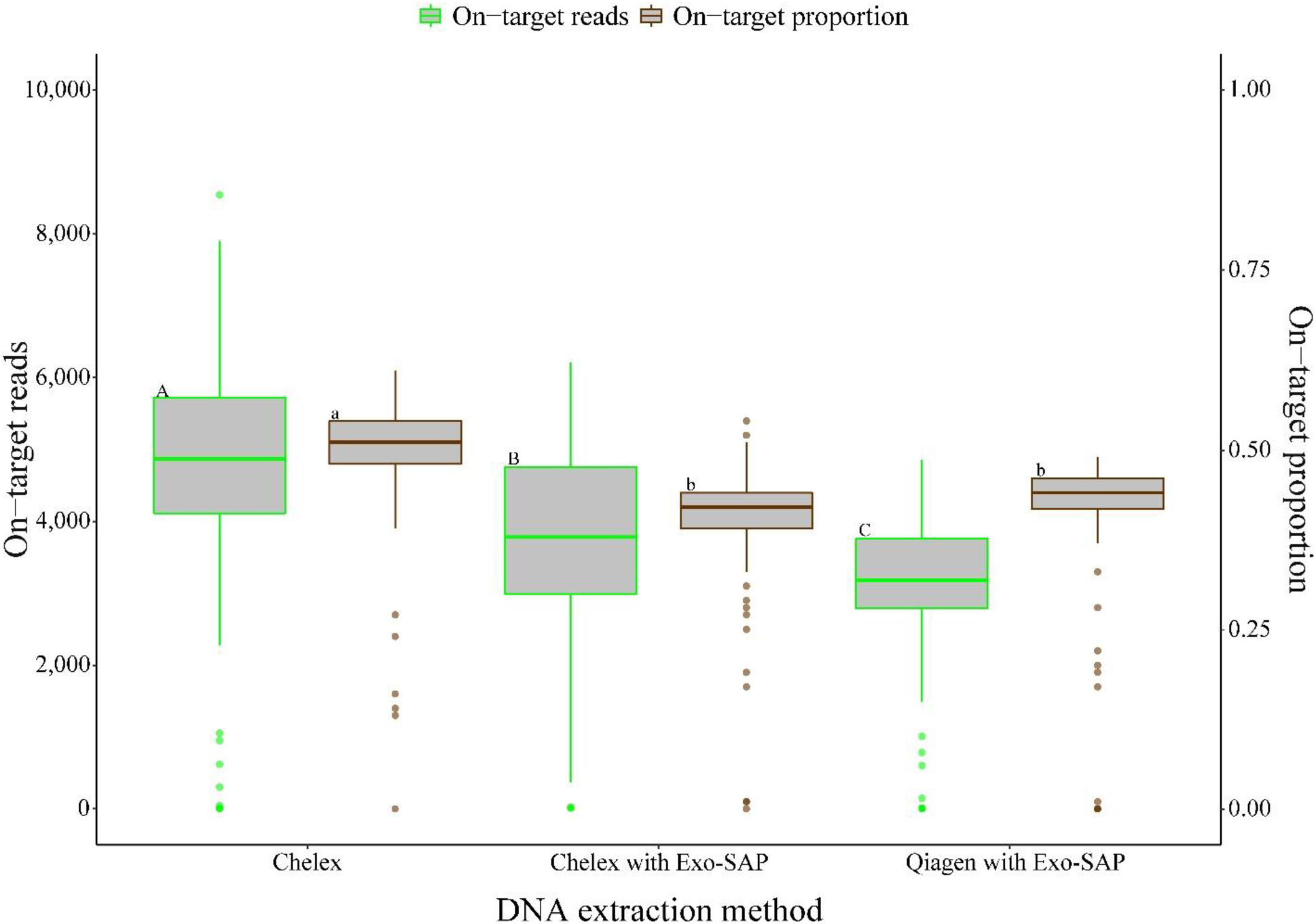
Number of on-target reads (green) and proportion of total reads on-target obtained from GT-seq libraries produced using DNAs extracted via Chelex, Chelex with Exo-SAP, and Qiagen with Exo-SAP. Significantly different groups denoted by letters on box.

A total of 4,063 genotypes across 406 loci (820 SNPs) could be used in comparisons between GT-seq data and those obtained from the original RAD study. Of these genotypes, 96.6% of calls were identical between methods, and modeled expectations of genotype concordance (residual sum of squares = 0.02) indicated that a concordance rate of 99.0% could be expected at a GT-seq read depth of 31 (Fig. 6).

**Figure 6.**
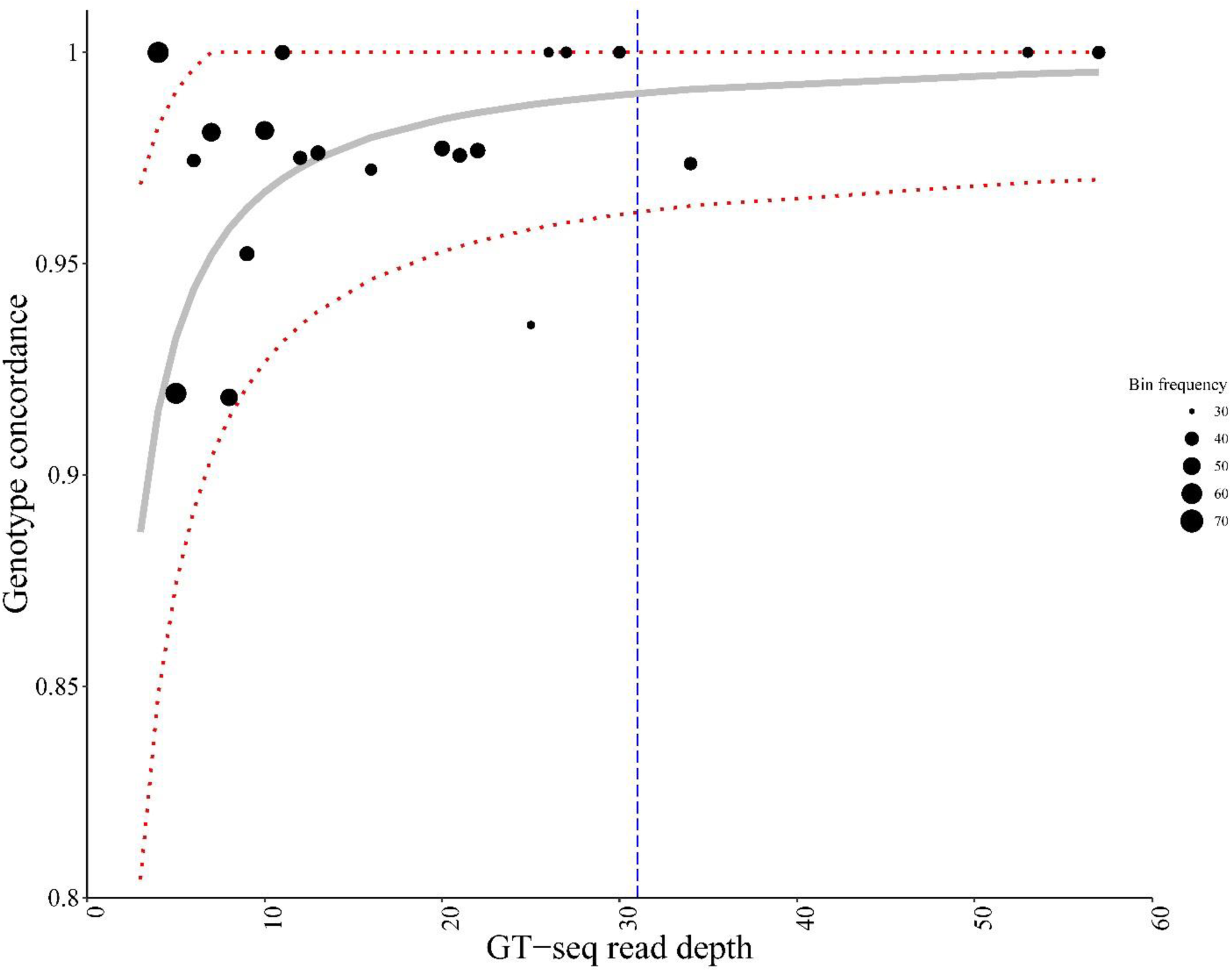
Modeled relationship between GT-seq read depth and genotype concordance between GT-seq and RADseq shown in gray (1.00-0.34/GT-seq read depth, rss = 0.02) with 95% confidence intervals in red. GT-seq read depth at which estimated genotype concordance equals 99% (96.2%-100%) represented by blue line. Black points display proportion of genotypes found identical between GT-seq and RADseq for GT-seq read depth bins with > 30 genotypes.

For a final proof of concept, a new sample of 570 walleye was sequenced using the current panel of 436 loci. After filtering, 551 individuals and 303 loci were retained with an average of 32.9 (SD = 29.1) reads per locus; 116 of the 303 loci exhibited an average coverage greater than the 31× target identified for 99.0% genotyping concordance (Fig. 7). The average percent of missing data was 6.4% (SD = 13.0%) across individuals and 30.0% (SD = 38.0%) across loci.

**Figure 7.**
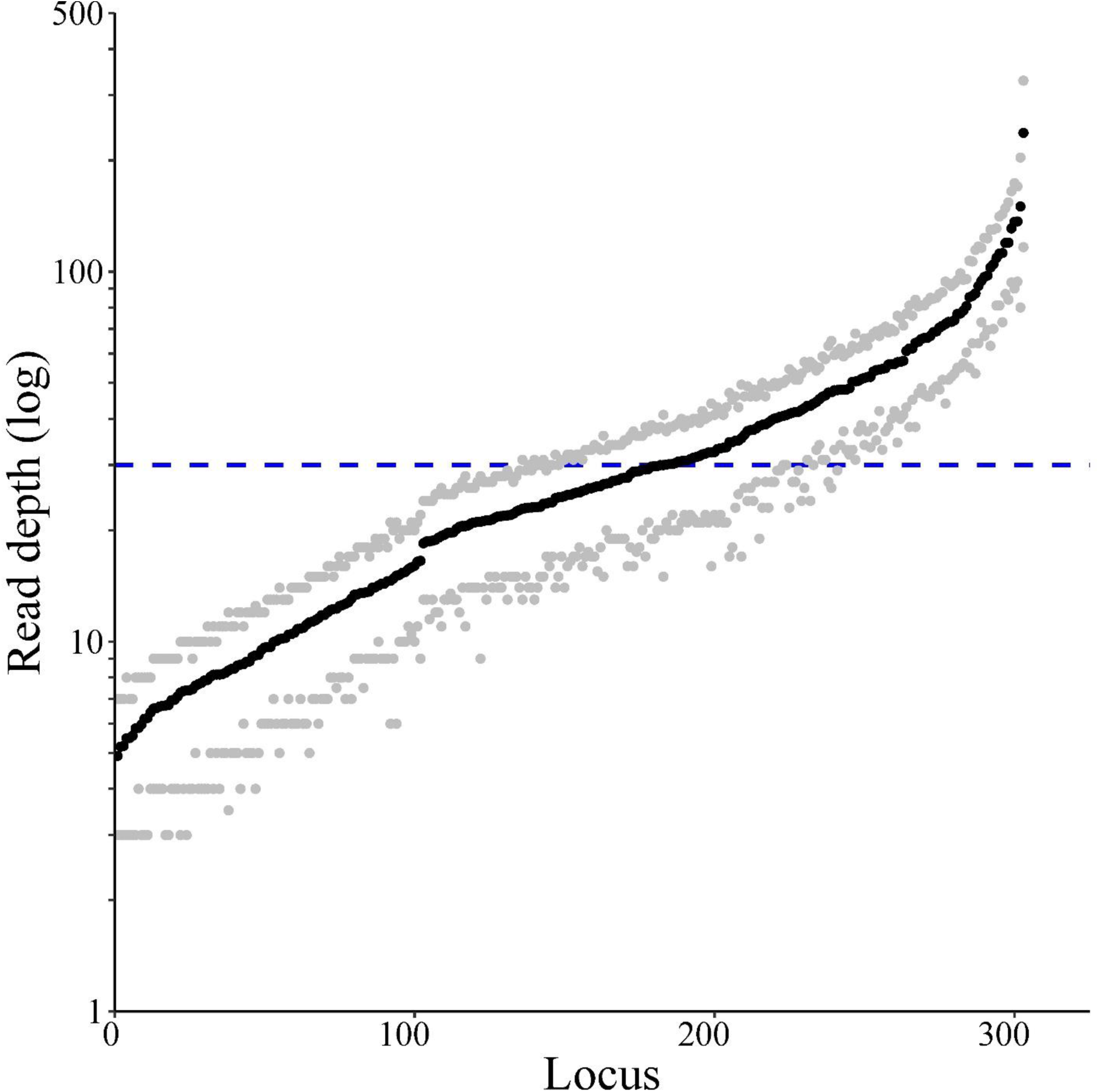
Variation in read depth among individuals at loci successfully genotyped after quality filtering (303 loci with < 30% missing data). Average read depth at each locus shown with black points, while gray points denote first and third quartile for each locus. Dotted blue line denotes target read depth of 30×. Data from 551 walleye sequenced using fully optimized panel. Average read depth among all loci is 33×.

## Discussion

GT-seq and other amplicon sequencing methods have tremendous potential for facilitating high-throughput genotyping in non-model organisms (Meek & Larson, 2019). The general steps for GT-seq panel development: SNP ascertainment, SNP selection, primer design, and panel optimization have been previously detailed (Baetscher et al., 2018; McKinney et al., 2019; Schmidt et al., 2019); however, the process of GT-seq panel development is not static. Here, we leverage our experiences developing a GT-seq panel for walleye with testing various aspects of the GT-seq methodological process to provide additional guidelines usable by other researchers to simplify panel construction and validation, particularly in non-model species. Our walleye panel has the necessary power to conduct GSI in a study system with highly variable degrees of genetic differentiation and perturbation by historical stocking, while also being capable of identifying PO and FS relationships within large populations. The robust performance of our panel was facilitated by exploring the upper limits of how many loci a GT-seq panel can target and the trade-offs between choosing loci for GSI versus parentage analysis. The information presented here will aid in the efficient creation of multipurpose GT-seq panels in organisms with little to no available genomic resources.

### Patterns of population structure: historical stocking influences GSI accuracy

The largest genetic differentiation in our data was observed between populations from Wisconsin and Minnesota; this structure was likely the result of recolonization from different refugia following the Wisconsin glaciation, which ended ~10,000 years ago. A range-wide analysis of walleye genetic structure using microsatellite loci produced similar patterns, with the most genetically independent populations found in northern Minnesota and Canada (Stepien et al., 2009). Additionally, we found that while populations in Minnesota displayed strong isolation on relatively small spatial scales, broad-scale patterns of isolation were less evident in Wisconsin. In particular, the Ceded Territory of Wisconsin, which included our Chippewa River and Wisconsin River reporting groups, displayed patchy and low genetic structure overall. It is likely that structure in this region has been compromised by stocking. Hammen and Sloss (2019), for instance, observed that several populations of walleye in the upper Chippewa were more genetically similar to populations in the upper Wisconsin than to other populations in the upper Chippewa, while nongame species in the Ceded Territory of Wisconsin displayed patterns of genetic divergence strictly associated with drainage basin boundaries (Westbrook, 2012). We also observed that four proximate populations spanning the Chippewa and Wisconsin River boundaries were nearly indistinguishable (Turtle Flambeau Flowage, Manitowish Lake, Kawaguesaga Lake, Big Arbor Vitae Lake). These populations are within 50 km of each other and are located near a state walleye hatchery in Woodruff, Wisconsin, that has historically used broodstock solely from the Wisconsin River drainage basin. It is therefore highly likely that the genetic similarity of these four populations is due to stocking. Several of the sampled populations from Minnesota also had poorly documented stocking histories yet they remained highly distinct. Genetic structure in Minnesota may have been less eroded if local, genetically similar sources were used, stocking was into larger, healthier resident populations, or stocking was less intense or ended a longer time ago.

Despite the challenges posed by low *F*_ST_ and evidence of supplemental stocking altering genetic structure in some populations, the SNPs discovered here provide greatly increased resolution for defining reporting units across the Midwestern, USA. Additionally, simulations suggested that a panel of several hundred loci would be highly capable of conducting individual-based GSI for most genetic units in the region. Given the regional complexity, however, improvements to accuracy could be made by further sampling areas that have shown heterogeneous signals of genetic structure (e.g., due to stocking). For example, increased sampling effort directed at the Chippewa and Wisconsin Rivers’ drainage basins could prove especially beneficial as analyzing populations in the lower reaches of each basin may provide a better understanding of signals of historical recolonization, while populations in the upper reaches (e.g., Ceded Territory of Wisconsin) could better define the effects stocking may have had. Additional samples could also serve as a holdout dataset, as suggested by Anderson (2010), to test the assignment accuracy of our panel.

### Tradeoffs associated with choosing loci based on differentiation versus diversity

We evaluated the tradeoffs associated with selecting SNPs based on differentiation or diversity and found that there was relatively little variation in GSI accuracies across panels. Markers selected based on differentiation have been shown to provide increased resolution for defining reporting groups in systems with low levels of genetic structure (Larson et al., 2014; McKinney et al., 2019). This approach has not, however, been applied to systems where stocking may be a major factor for reduced levels of population structure, such as in upper Midwestern, USA, walleye. Interestingly, we found that assignment accuracies with our smaller panels was relatively similar to accuracies obtained using ~30,000 SNPs discovered with RAD-seq (data not shown). This suggests that assignment accuracy in our system may be limited more by biological realities associated with human-mediated gene flow than by the power of our genetic markers. Further increases in assignment accuracy are therefore likely to be realized through sampling of additional populations and a more refined understanding of population history as opposed to genotyping additional markers.

Conversely, we found that FPRs for assigning kin relationships were highly variable among panels, with the microhaplotype diversity-based panel displaying the lowest FPRs by several orders of magnitude for each kin relationship (Table 3). This contrast in inter-panel variation between GSI and kinship simulations is reflective of the variation in information content of each panel (Fig. S1), and supports previous findings that while microhaplotype information provided added benefit to both applications, the greatest increase in assignment accuracy will likely be for kinship analysis (Baetscher et al., 2018; McKinney, Seeb, et al., 2017). When attempting to target microhaplotype loci via GT-seq, attention should be given to the number of SNPs one aims to genotype within a locus, as attempting to include loci with too many SNPs may result in targeting repetitive regions that fail to amplify properly in a multiplex. The expected maximum number of alleles per locus and the degree to which loci with large numbers of alleles perturbs primer design will likely vary among taxa. We chose a cutoff of 10 alleles per locus as this appeared to be a natural break point in the allele distribution for walleye; we suggest that researchers investigate this in their system and come up with a logical cutoff prior to selecting loci. Finally, while our results suggested this panel could facilitate HS identification in small systems, performing this task in large systems would likely require more loci. Our tests of panel implementation suggest this could be achievable by combining PCR products from several panels within individuals prior to barcoding.

### Optimizing primer design and removing overamplifying loci

The main objective of GT-seq primer development is to produce a single pool of primer pairs that will amplify uniformly, while retaining as many loci as possible. To achieve this, it is important to minimize heterogeneity of primer and product characteristics (e.g., primer size, product size) and to understand that the highly multiplexed PCR required by GT-seq can be complicated by hairpin- and inter-primer hybridization artifacts. To best control PCR artifacts, it is important to avoid developing primers with complimentary regions (e.g., complimentary 3’ regions and self-complementarity) and apply conservative thresholds to the upper T_m_ of primer design parameters (Rychlik, 1993). Incorporating loci with multiple SNPs can lead to further difficulties when the ideal priming region also contains a SNP. We found that, while degenerate primers could be successfully amplified in a multiplex, they were culled during optimization at a higher rate than non-degenerate primers. Further performance benefits could be gained from examining DNA template quality beyond just the availability of priming regions, as shown by Benita et al. (2003) who found regionalized GC content of template DNA to be a predictor of PCR success. This was supported by our data, as loci removed from the panel during optimization displayed significantly higher GC content in the amplicon and primer. Finally, while GT-seq primers can theoretically be designed for a range of amplicon sizes, we suggest that researchers design panels targeting similarly sized products that can be sequenced using PE150 technology. Panels containing similarly sized and relatively short amplicons should reduce variation in amplification rates (Baetscher et al., 2018) and ensure that genotyping is robust to variation in sample quality. Moreover, PE150 sequencing is common to benchtop and core facility sequencing platforms, such as Illumina® MiSeq and HiSeq.

In exploring the upper limits of how many loci a GT-seq panel can target, we found that the number of amplicons reliably genotyped in a single pool is highly dependent on variable rates of amplification among primer pairs during PCR and, to a lesser extent, the degree of primer specificity. Despite efforts to limit primer inter-hybridization through diligent primer design, the presence of overamplifying loci is likely inevitable during early phases of panel development (see also McKinney et al. 2019). We found it best to focus primarily on the uniformity of amplification within the primer pool in early optimization steps, by removing primer pairs found to overamplify. Although achieving perfect uniformity is challenging, application of strict cutoffs during initial optimization steps likely results in a final panel that is less influenced by overamplification, thereby increasing the upper limit of GT-seq performance. The importance of this was illustrated by prop_reads_T10 reducing from 85.0% of all primer reads to 36.6% after optimization. Likewise, on-target rates were greatly improved by addressing overamplification, as demonstrated by the on-target proportion of reads increasing from 24.9% to 91.7% by the third test.

### Further optimization of the GT-seq protocol

Although there may be an upper as-yet-unidentified limit in the number of primers that can be included in a single primer pool, we found that the total number of loci targeted can be increased by PCR amplifying multiple primer pools separately on a sample and pooling PCR products within individuals prior to barcoding. This approach could be used to genotype multiple complementary or even independent GT-seq panels using the same primer tail systems at a small cost increase compared to genotyping a single panel, as the most expensive steps in the GT-seq protocol (e.g., DNA normalization) are only conducted once (Campbell et al., 2015). Combining multiple panels could facilitate genotyping of > 1,000 loci rather than a few hundred, providing greatly increased power for kinship analysis and GSI (Baetscher et al., 2018; McKinney, Seeb, et al., 2017). Additionally, further optimization of individual panels could be conducted by manipulating the initial concentrations of primer pairs based on observed panel performance, reducing the concentration of loci that appear to overamplify. While this process would be cumbersome to perform by hand, a liquid handling robot could enable a researcher to fine-tune the performance of existing and new panels alike, thereby enhancing efficiency.

DNA extraction can comprise a large portion of the total cost of genetic analysis, especially for relatively affordable approaches such as GT-seq, in terms of finances and time. Extractions using chelating beads provided a cost-effective alternative to more expensive salting-out approaches, such as Qiagen DNeasy kits. Chelating extractions, however, can also produce lower quality DNA and may include suspended impurities (Singh et al., 2018). Campbell et al. (2015) did show that GT-seq can be conducted using DNA from chelating extractions but did not directly compare results using multiple extraction protocols. Here, we directly showed that cost-effective chelating extractions can produce equally high quality, if not superior, sequence data compared to more expensive methods. Although consideration should be given to the quality of tissue samples, the chelating approach appears to be a viable approach for reducing per-sample costs with GT-seq. It is important to be aware that proper laboratory technique is essential when using this method, however, as chelating beads will inhibit PCR and greatly reduce library product yields. This may be especially problematic when using a liquid handling robot that is unable to visually detect chelating beads. Therefore, we suggest researchers carefully pipette the DNA-containing supernatant from chelating resin extractions by hand into a secondary container (e.g., 96-well PCR plate) before aliquoting DNA with a robot. Finally, we found that the
ExoSAP procedure included in the original GT-seq protocol did not produce higher quality data and was not necessary for our purposes; removing this step from the protocol will further reduce GT-seq costs and time commitment.

### Suggestions for designing GT-seq studies and conclusions

A major consideration when designing a GT-seq panel is deciding how large of an ascertainment dataset is necessary. We constructed a comprehensive ascertainment set with RAD-seq, which was expensive and resource intensive. Despite this, we found that the panel chosen based on diversity produced similar results to the panel chosen based on differentiation. In our case, we believe that a smaller ascertainment set of ~96 individuals sampled from across the same geographic range may have resulted in a panel of relatively similar quality. Smaller ascertainment datasets are likely sufficient when the main applications of a given GT-seq panel are kinship analysis and GSI of highly diverged populations; however, when designing GT-seq panels to differentiate closely related populations (e.g. Chinook salmon *Oncorhynchus tshawytscha* in western Alaska), accurate characterization of ascertainment populations is vital (Larson et al., 2014; McKinney et al., 2019).

Another major consideration when conducting GT-seq analysis is deciding how deep to sequence individuals. We found that a read depth of 31× could be expected to produce genotypes that were 99% concordant with those derived from RADseq. Read depths were, however, highly variable across loci; we only retained 303 of the 436 loci in our panel when we genotyped 536 individuals at an average depth of 33×. We also found that a large and variable proportion of reads can be discarded prior to genotyping. Therefore, we suggest that researchers target an average depth of at least 100× to ensure that most loci in the panel can be genotyped and that all acquired genotypes are highly reliable. At this level of coverage, researchers could genotype ~500 individuals with a panel of 500 loci on a single MiSeq lane (~25 million reads) and ~8,000 individuals on a HiSeq lane (~400 million reads). It is possible this level of coverage is not necessary for some applications, such as GSI, but we strongly suggest obtaining high coverage for more sensitive applications that require high genotyping accuracy, such as kinship analysis.

Finally, researchers conducting GT-seq must consider trade-offs associated with different genotyping approaches. The two main approaches we are aware of are: (1) in-silico probe-based methods that use pattern matching to genotype specific alleles (Campbell et al., 2015; McKinney et al., 2019) and (2) alignment-based methods that call all polymorphisms in a given amplicon (Baetscher et al., 2019). A major advantage of probe-based methods is that databases of probes can be shared among laboratories, facilitating standardization. It is difficult, however, to discover new variation with these methods, whereas alignment-based methods discover new variation by default. We suggest a hybrid approach, where researchers periodically use alignment-based approaches to discover new variation and add this variation to a probe database that forms the basis of genotyping and standardizing genotyping among laboratories.

GT-seq is a powerful addition to the molecular ecologist’s toolkit that facilitates rapid, accurate, and cost-effective genetic analysis. Yet, creating a GT-seq panel is non-trivial, and there are many considerations for maximizing the utility of this approach. We found that the greatest challenge when designing our GT-seq panel was locus-specific overamplification, and we suggest that researchers remove these loci liberally. We also found that chelating extractions without an ExoSAP step produce high-quality results, providing a lower-cost alternative to salting-out extractions. Additionally, we showed that combining multiplex PCR products from multiple panels prior to barcoding can ensure additional, potentially important, loci can be genotyped with only a moderate cost increase. Finally, we found that a relatively substantial proportion of sequencing reads are lost before genotyping, and we suggest researchers target higher sequencing coverage (100×) than may apparently be necessary to ensure that GT-seq datasets are robust across loci. The GT-seq approach promises to be a mainstay of population genetics for the foreseeable future, and the guidelines and suggestions outlined here may help increase the effective use of this powerful method.

## Acknowledgements

We thank field crews from the Wisconsin and Minnesota Departments of Natural Resources for collecting tissue samples. We also thank Keith Turnquist from UW-Stevens Point for assistance with laboratory analysis and Ana Ramón-Laca for advice regarding primer development. This study was funded by the United States Fish and Wildlife Service, Federal Aid in Sportfish Restoration program and the Wisconsin Department of Natural Resources. Any use of trade, firm, or product names is for descriptive purposes only and does not imply endorsement by the U.S. Government.

## Data accessibility

Raw data for the RADseq and GT-seq data obtained in this study was deposited to the NCBI sequence read archive (SUB####) and VCF files of genotypes are available on DRYAD (DOI: PENDING). Python and R scripts used in the statistical analysis pipeline are available at GIT

## Author contributions

WL, GS, KG, and LM designed the study with input from MB. Data analyses were conducted by MB with assistance from GM. Laboratory analysis was conducted by MB, KG, and LS. All authors contributed to the writing of the manuscript.

## Supplementary materials

**Table S1.** Pairwise *F*_ST_ estimates for all sampled walleye *Sander vitreus* populations (sites numbered according to Table 1 and Fig. 1 A). Estimates produced in arlequin v3.5.2.

**Table S2.** Summary statistics for 20,597 SNPs retained through initial filtering based on maximum missingness rates of < 20% and HDplot cutoffs of H > 0.5 and −7 < D < 7. Columns include a locus tag (CHROM), position of SNP within locus (Reid et al.), a unique SNP value (ID), reference (REF) and alternate (Keenan et al.) SNP alleles, global *F*_IS_ (Willi et al.), single SNP *F*_ST_ (Smith et al.), expected microhaplotype heterozygosity (mhap_*H*_E_), and number of alleles per locus tag (n_alleles). Diversity statistics estimated in diveRsity v1.9.90 (global *F*_IS_ and single SNP *F*_ST_) and adegenet v2.1.1 (single locus *H*_E_, number of alleles).

**Table S3.** Summary matrix of 100% simulations (reps = 1,000, mixsize = 200) for each sampled population retained through filtering, performed using the *F*_ST_600_ panel. Each row represents a simulation for the listed population name. Each column within a row represents the proportion of individuals assigned to the population denoted at the top of the column. Unassigned individuals (< 70% probability of origin from a given population) accounted for in last column.

**Table S4.** Summary matrix of 100% simulations (reps = 1,000, mixsize = 200) for each sampled population retained through filtering steps, performed using the Composite__600_ panel. Each row represents a simulation for the listed population name. Each column within a row represents the proportion of individuals assigned to the population denoted at the top of the column. Unassigned individuals (< 70% probability of origin from a given population) are accounted for in the last column.

**Table S5.** Summary matrix of 100% simulations (reps = 1,000, mixsize = 200) for each sampled population retained through filtering steps, performed using the Diversity__600_ panel. Each row represents a simulation for the listed population name. Each column within a row represents the proportion of individuals assigned to the population denoted at the top of the column. Unassigned individuals (< 70% probability of origin from a given population) are accounted for in the last column.

---

**Figure S1.**
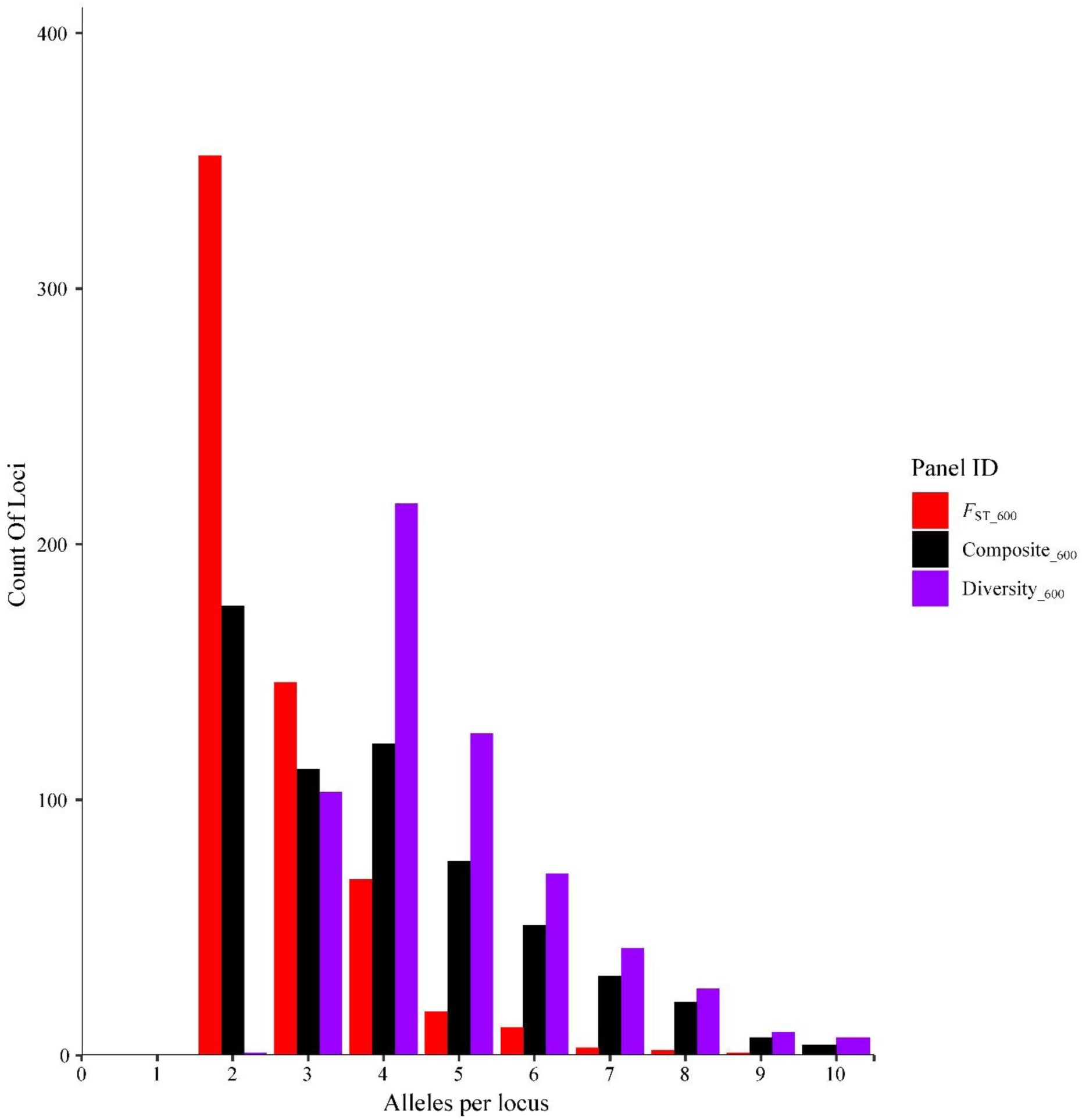
Frequency distribution of number of alleles among 600 loci tested in each panel.

